# Diversification and collapse of a telomere elongation mechanism

**DOI:** 10.1101/445429

**Authors:** Bastien Saint-Leandre, Son C. Nguyen, Mia T. Levine

**Affiliations:** Department of Biology, University of Pennsylvania, Philadelphia, PA, 19104.; Epigenetics Institute, University of Pennsylvania, Philadelphia, PA, 19104.; Department of Genetics, Perelman School of Medicine, University of Pennsylvania, Philadelphia, PA, 19104.

**Keywords:** telomere, retrotransposon, intra-genomic conflict, Drosophila

## Abstract

In virtually all eukaryotes, telomerase counteracts chromosome erosion by adding repetitive sequence to terminal ends. *Drosophila melanogaster* instead relies on specialized retrotransposons that insert preferentially at telomeres. This exchange of goods between host and mobile element—wherein the mobile element provides an essential genome service and the host provides a hospitable niche for mobile element propagation—has been called a ‘genomic symbiosis’. However, these telomere-specialized, ‘jockey’ family elements may actually evolve to selfishly over-replicate in the genomes that they ostensibly serve. Under this intra-genomic conflict model, we expect rapid diversification of telomere-specialized retrotransposon lineages and possibly, the breakdown of this tenuous relationship. Here we report data consistent with both predictions. Searching the raw reads of the 15-million-year-old ‘melanogaster species group’, we generated *de novo* jockey retrotransposon consensus sequences and used phylogenetic tree-building to delineate four distinct telomere-associated lineages. Recurrent gains, losses, and replacements account for this striking retrotransposon lineage diversity. Moreover, an ancestrally telomere-specialized element has ‘escaped,’ residing now throughout the genome of *D. rhopaloa.* In *D. biarmipes,* telomere-specialized elements have disappeared completely. *De novo* assembly of long-reads and cytogenetics confirmed this species-specific collapse of retrotransposon-dependent telomere elongation. Instead, telomere-restricted satellite DNA and DNA transposon fragments occupy its terminal ends. We infer that *D. biarmipes* relies instead on a recombination-based mechanism conserved from yeast to flies to humans. Combined with previous reports of adaptive evolution at host proteins that regulate telomere length, telomere-associated retrotransposon diversification and disappearance offer compelling evidence that intra-genomic conflict shapes Drosophila telomere evolution.

## Introduction

Transposable elements infest eukaryotic genomes, ever-evolving to increase in copy number over time (Feschotte and Pritham 2007; Beauregard et al. 2008). These ‘selfish elements’ compromise host fitness by disrupting host genes (Hancks and Kazazian 2012), nucleating local epigenetic silencing (Slotkin and Martienssen 2007; Lee and Karpen 2017), and triggering catastrophic recombination between non-homologous genomic regions (Langley et al. 1988; Beck et al. 2011). Transposable elements (‘TEs’) also provide raw material for genome adaptation (Jangam et al. 2017): across eukaryotes, host genomes re-purpose TE-derived sequence for basic cellular and developmental processes, from immune response (van de Lagemaat et al. 2003) to placental development (Lynch et al. 2015) to programmed genome rearrangements (Cheng et al. 2010; Cheng et al. 2016). These diverse ‘molecular domestication’ events share a common feature—the degeneration or deletion of the TE’s capacity to propagate. Consequently, the TE-derived sequence resides permanently at a single genome location, just like any other host gene sequence. Inability to selfishly over-replicate resolves prior conflict of interest between the host and the TE (Jangam et al. 2017). However, not all adaptive molecular domestication events necessitate TE immobilization; in rare cases, essential host functions rely on retention of the mobilization machinery that promotes recurrent TE insertions into host DNA. The non-canonical telomere elongation mechanism of Drosophila is exemplary (Pardue and DeBaryshe 2003; Casacuberta 2017).

In virtually all eukaryotes beyond Drosophila, telomerase-added DNA repeats (Greider and Blackburn 1989; Zakian 1989; Blackburn 1991; Zakian 1996) counteract the ‘end-replication problem’ that otherwise erodes unique DNA sequence at chromosome termini (Watson 1972). However, telomeres of select fungi (Starnes et al. 2012), algae (Higashiyama et al. 1997), moths (Osanai-Futahashi and Fujiwara 2011), crustaceans (Gladyshev and Arkhipova 2007), and DNA repair-deficient mammalian cells (Morrish et al. 2007) encode not only telomerase- added repeat elements but also TEs that insert preferentially at chromosome termini. These telomeric mobile elements are typically derived from a single class of TE—the non-long terminal repeat (“non-LTR”) retrotransposons (Beck et al. 2011), which mobilize via reverse transcription and insertion into new genomic locations. The most extreme example of this co-option is found in *Drosophila melanogaster,* whose telomeres harbor no telomerase-added repeats. In fact, the 220 million-year-old ‘true fly’ insect Order, Diptera, completely lacks the genes encoding the telomerase holoenzyme (Pardue and DeBaryshe 2003; Casacuberta 2017). Instead, *D. melanogaster* harbors three highly specialized retrotransposons—HeT-A, TART, and TAHRE—charged with the important job of preserving distal, unique sequence (Pardue and DeBaryshe 2011). These telomere-specialized retrotransposons represent a monophyletic clade within the larger ‘jockey’ element family (Villasante et al. 2007), whose members are more typically found along chromosome arms (Xie et al. 2013). The telomere-specialized jockey subclade, in contrast, rarely inserts outside the telomere (Pardue and DeBaryshe 2003; Berloco et al. 2005; Pardue and DeBaryshe 2011).

This molecular domestication of still-mobile retrotransposons into an essential genome function is often referred to as a genomic ‘symbiosis’ (Pardue and DeBaryshe 2008). Evidence for such a mutualism is compelling. The elements that maintain telomere ends in *D. melanogaster* comprise a monophyletic clade ostensibly specialized to replicate only at chromosome ends (Pardue and DeBaryshe 2008). Elements from this jockey clade also appear at terminal ends in distant Drosophila species, consistent with a single domestication event >40 million years ago followed by faithful vertical transmission (Casacuberta and Pardue 2003; Villasante et al. 2007). Moreover, no other mobile element family appears at terminal ends, consistent with host-dependent molecular mechanisms that block ‘generalist’ TEs from invading this vital genome region (Mason and Biessmann 1995; Mason et al. 2016). These data suggest that retrotransposon-mediated chromosome elongation represents a long-term, conserved relationship between a host genome and its domesticated, but still-mobile, retrotransposons.

This picture of cooperativity was complicated by the discovery that the telomere-associated subclade of jockey elements evolves rapidly across Drosophila. Leveraging 12 Drosophila genomes that span 40 million years of evolution, Villasante, Abad, and colleagues detected at least one jockey-like, candidate telomeric retrotransposon in all 12 species (Villasante et al. 2007). These data implicated a single evolutionary event in a common ancestral sequence that conferred telomere-specificity. However, these elements represent distinct phylogenetic lineages rather than species-specific versions of the HeT-A, TART, and TAHRE elements well-studied in *D. melanogaster.* For example, *D. melanogaster* and its close relatives encode HeT-A, TART, and TAHRE, while the 20 million year-diverged *D. ananassae* encodes a single, phylogenetically distinct candidate retrotransposon lineage, ‘TR2’. The 30 million-year diverged *D. pseudoobscura* species encodes yet other phylogenetically distinct lineages within this jockey subclade. This expansive evolutionary lens revealed a previously unappreciated, dynamic evolutionary history of these jockey subclade retrotransposons. However, the large evolutionary distances between species left fine scale dynamics unknown and telomere-specific localization was not explored (Villasante et al. 2007). The evolutionary origin(s), ages, between-species differences, and genome locations of these candidate length regulators remain obscure. Elucidating the evolutionary history of these elements is essential to address the possibility that this molecular domestication event is less a stable, long-term genomic symbiosis and instead a tenuous relationship between the domesticator and the domesticated.

Our recent report that pervasive positive selection shapes Drosophila telomere proteins— especially genes that regulate chromosome length (Lee et al. 2017)—raises the intriguing possibility that telomeric retrotransposons recurrently evolve to escape host control. Under this model, domesticated retrotransposons evolve to increase copy number either by excessively lengthening chromosome ends or by invading non-telomeric locations. Host dependency on still-mobile TEs to perform an essential genome function may, in fact, be an unstable telomere elongation strategy. To evaluate this possibility, we set out to resolve the finer scale evolutionary history of telomere-specialized elements. Specifically, we investigated the ‘melanogaster species group’, which captures an informative three-15 million years of evolution (Chen et al. 2014). This clade spans the distance between HeT-A-, TART-, and TARHE- encoding *D. melanogaster* and its close relatives and *D. ananassae,* which encodes the distant TR2 lineage not yet localized to telomeres (Danilevskaya et al. 1998; Casacuberta and Pardue 2002; Villasante et al. 2007). We probed the raw reads of melanogaster species group genomes and used *de novo* consensus building and experimental validation to define young and ancient telomere-associated retrotransposons from this jockey subclade. We uncovered diversification via gain and loss, both with and without replacement, of telomere-associated, retrotransposon lineages across three to 15 million years of evolution. We also observe dramatic differences in telomere-specialized retrotransposon copy number across these closely related species, consistent with chromosome length evolving under minimal host constraint. In the most extreme case, we identified a species that has undergone wholesale loss of active, telomere-specialized retrotransposons. Collapse of this telomere elongation mechanism highlights the potential hazards of depending on still-mobile elements to perform an essential genome service.

## Results

### Candidate telomere-specialized retrotransposons identified in the melanogaster species group

*D. melanogaster* encodes telomere-specialized, non-LTR retrotransposons that increase in copy number by a “copy and paste” mechanism (Pardue and DeBaryshe 2011). Transcripts encoded by these elements are localized, reverse transcribed, and integrated at the terminal nucleotides of chromosome ends, resulting in stereotypical head-to-tail arrays (Pardue and DeBaryshe 2003). Full length, autonomous retrotransposons typically contain two open reading frames (ORFs) between the variable 5’ and 3’ UTRs. ORF1 encodes an RNA binding domain (‘Gag’) and ORF2 encodes a reverse transcriptase domain (“RT”) and an endonuclease domain (“EN”, Fig. S1). The telomere-specialized HeT-A (ORF1 only), TART, and TAHRE elements form a monophyletic clade within the jockey family of non-LTR retrotransposons (Villasante et al. 2007).

To elucidate the fine scale evolutionary history of telomere-specialized elements in Drosophila, we searched for jockey-like, telomere-specialized retrotransposons across lineages that span three-15 million years of evolution (Drosophila 12 Genomes et al. 2007; Chen et al. 2014). *D. melanogaster* and its close relatives (the ‘melanogaster subgroup,’ including *D. melanogaster, D. simulans, D. sechellia, D. yakuba*) share the well-studied retrotransposon lineages HeT-A, TART, and TAHRE (Danilevskaya et al. 1998; Casacuberta and Pardue 2002; Berloco et al. 2005; Villasante et al. 2007) and so offer limited traction for capturing evolutionary turnover events of major telomeric retrotransposon lineages. Beyond these close relatives, previous work defined a candidate telomeric retrotransposon from the jockey family in the 15-million year-diverged *D. ananassae* (Villasante et al. 2007). This element, however, represents a lineage distinct from HeT-A, TART, and TAHRE and has yet to be validated cytogenetically as telomere-specialized. To gain both sufficient divergence to reveal telomeric retrotransposon turnover and sufficient resolution to estimate retrotransposon lineage birth and death, we focused on an intermediate evolutionary distance from *D. melanogaster* by including other members of the ‘melanogaster species group’. Five focal species—*D. rhopaloa, D. biarmipes, D. takahsashii, D. elegans* and *D. ficusphila* — span 10 million years of divergence and share a common ancestor with *D. melanogaster* <15 million years ago (Chen et al. 2014). The genomes of *D. melanogaster* and its three close relatives served as positive controls for our *de novo* identification of retrotransposons phylogenetically related to the jockey subclade specialized at telomeres. *D. ananassae* served as an outgroup.

We developed a custom pipeline to discover jockey family elements ancestrally related to previously defined lineages that maintain chromosome ends (Fig. S2). Briefly, we conducted tBLASTn searches against raw reads from each of the ten species using a query that included both fully and minimally validated telomere-specialized retrotransposons from across Drosophila (Table S1). We also included the reference non-telomeric jockey element (from *D. melanogaster)* defined in Repbase (www.girinst.org/Repbase/, Table S1). To facilitate alignment across distant lineages, we restricted our search to the Gag domain of ORF1 and the RT of ORF2 (Fig. S1). These 63 Gag and 48 RT domains span 40 million years of Drosophila evolution (Table S1). A detailed description of our pipeline of iterative BLAST searches to raw reads, *de novo* consensus building, and phylogenetic tree-assisted sorting can be found in the Materials and Methods. This pipeline (Fig. S2) generated a refined list of consensus sequences that included previously described telomere-specialized elements from *D. melanogaster* and its close relatives (our “positive controls”) as well as jockey family elements outside the specialized telomeric subclade (Table S2). These latter elements indicated that our search was exhaustive—we effectively overshot the jockey subclade associated with telomeres for all species. We built Bayesian phylogenetic trees based on the Gag and RT domains of all final consensus sequences (Fig. 1 and Fig. S3). Our Gag-based trees revealed that the 15 consensus sequences form a well-supported, monophyletic subclade within the jockey family but distinct from 18 generalist jockey element consensus sequences that too form a distinct, well-supported monophyletic clade (gray, Fig 1A). The candidate telomeric Gag consensus sequences form four distinct lineages—TAHRE, TART, TR2, and a previously undefined lineage that we named ‘TARTAHRE’ for its labile phylogenetic position between TART and TAHRE. The TR2 lineage shares a more recent common ancestor with jockey elements compared to the TART/TAHRE/TARTHARE clade. Moreover, our trees support previous inferences that HeT-A Gags are phylogenetically indistinguishable from TAHRE Gags ((Villasante et al. 2007), Fig. S3)—HeT-A encodes its own 5’UTR, Gag, and 3’UTR (Pardue and DeBaryshe 2003), yet its phylogenetic position reveals that this single-domain element is effectively a TAHRE element missing an RT domain. Henceforth, we refer to this single lineage as TAHRE. The RT domain-based tree (Fig. 1B) revealed strikingly similar topology. We were surprised to recover complete consensus sequences of candidate telomere-specialized elements from only seven species of the 10 species. Our pipeline detected only a partial TAHRE Gag in *D. takahashii* (Table S2) and no evidence at all of the telomere-associated, jockey subclade elements in its close relative, *D. biarmipes* (see below). We explore below the possibility that *D. biarmipes* telomeres are functionally distinct from other sampled species.

**Figure 1.**
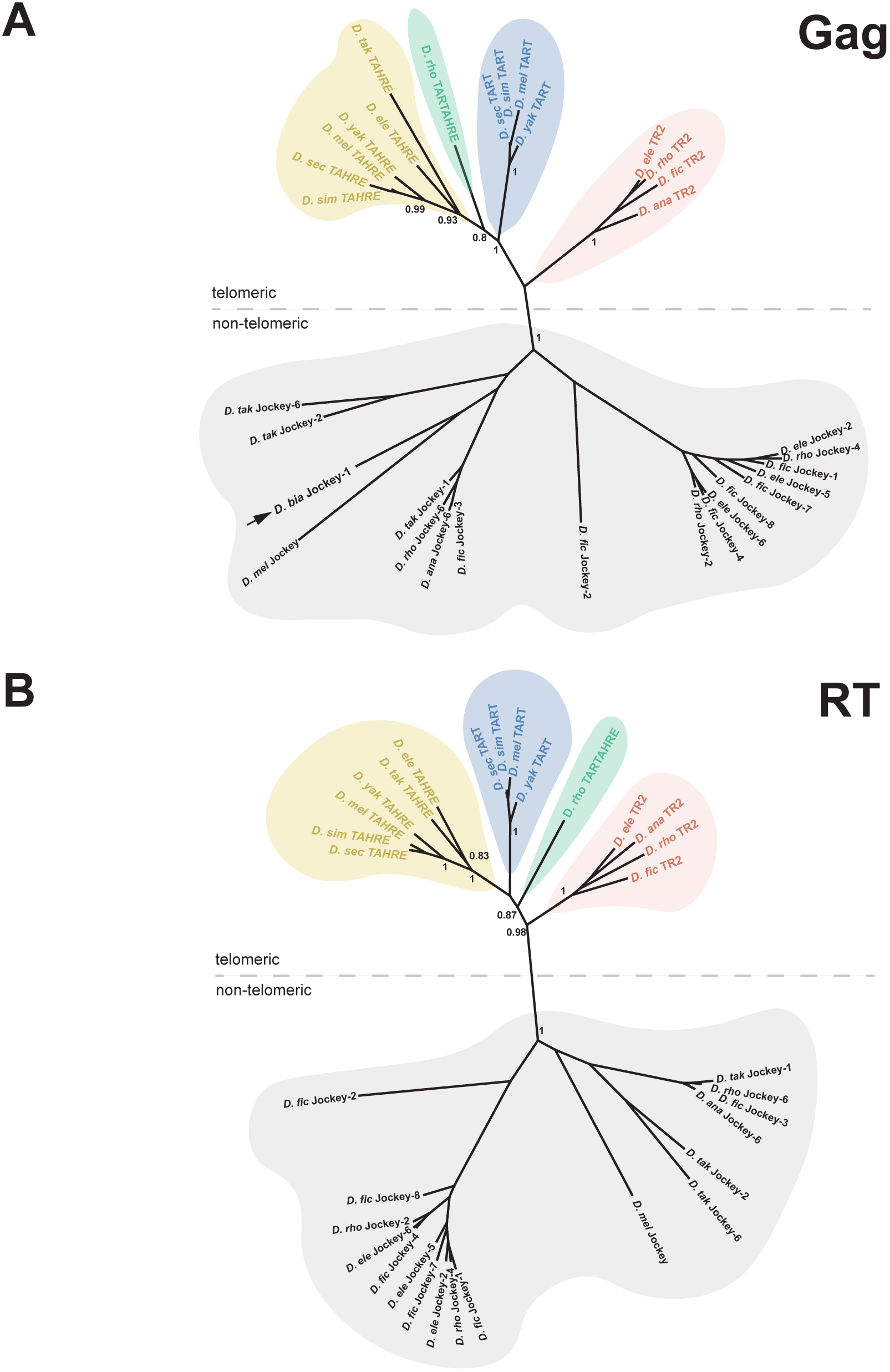
Phylogenetic relationships among previously and newly defined jockey subclade elements. Unrooted phylogenetic trees built from Gag domain **(A)** and RT domain **(B)** consensus sequences. Node support values are posterior probabilities generated by MrBayes. Gray designates jockey elements that passed the final pipeline filter but are distantly related to the telomere-specialized subclade. Various colors delineate candidate telomere-specialized elements along with previously characterized elements that form monophyletic clades. Only the D. rhopaloa-restricted element, TARTARHE (green), ‘migrates’ between TAHRE and TART clades across the two trees and may represent a chimera of the two lineages. The black arrow corresponds to the closest D. biarmipes jockey family element to the telomere-specialized subclade.

### PCR and cytogenetic validation of computationally predicted telomeric retrotransposons

Despite our search being inherently conservative—our consensus-building approach may average out sub-lineages within major named retrotransposon lineages—we uncovered striking diversity across only 15 million years of Drosophila evolution. By virtue of phylogenetic relatedness, we predict that the jockey subclade retrotransposons uncovered by our pipeline specialize at chromosome ends in their respective host species. Testing this prediction first requires molecular biology to confirm that 1) these *in silico-generated* consensus sequences represent actual elements in the targeted genomes and 2) the elements are arrayed in characteristic head-to-tail orientation that arises from exclusive end-integration of the poly-adenylated 3’ end (Pardue and DeBaryshe 2008; Pardue and DeBaryshe 2011). Using Sanger sequencing of PCR products amplified from genomic DNA, we discovered that the *in silico* sequences represent true DNA elements in their host genomes (Fig. 2A).

**Figure 2.**
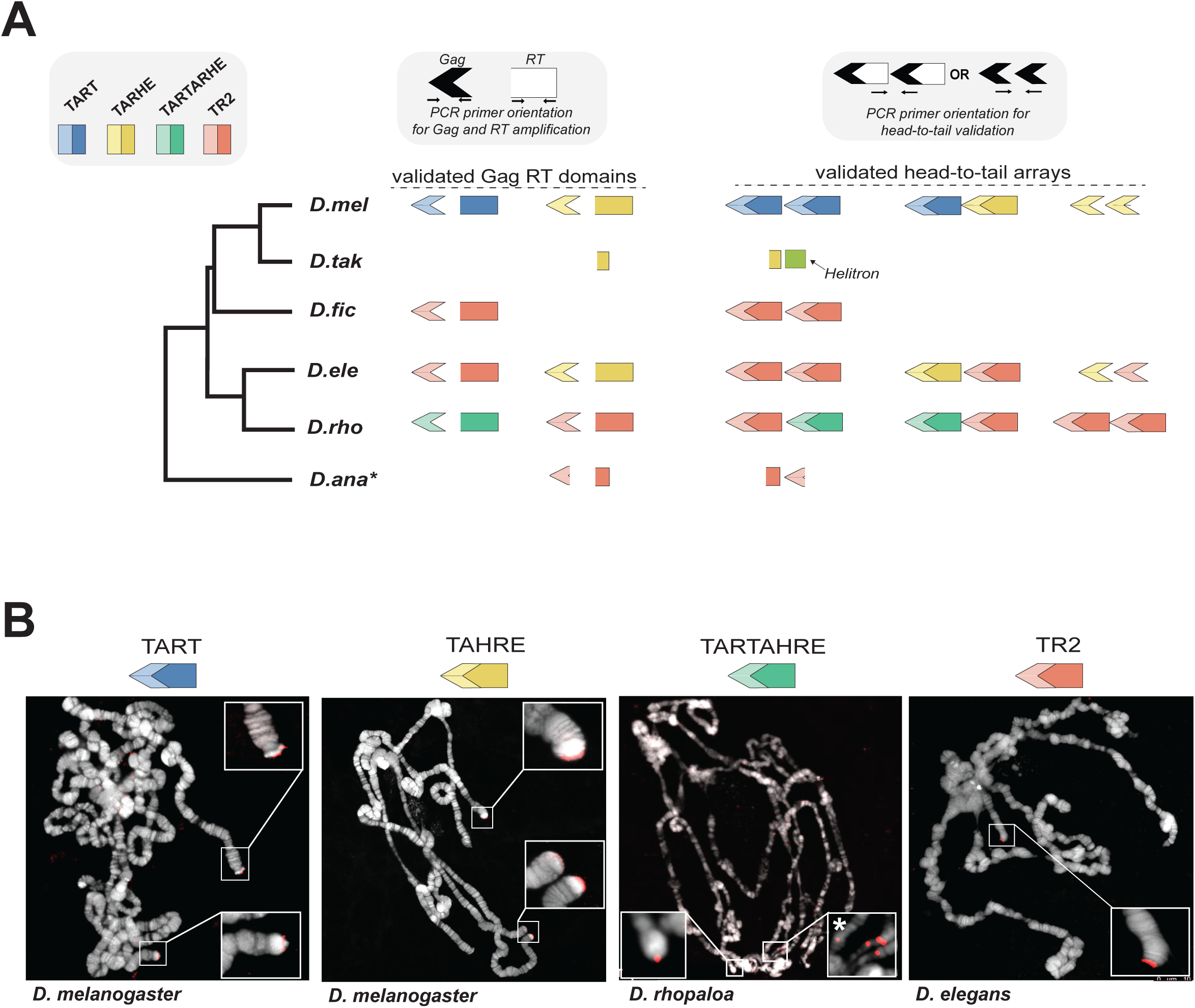
PCR- and cytology-based validation of in silico predicted, telomere-specialized elements. **(A)** Cartoon representation of our PCR-based validation of in silico – predicted Gag and RT domains and head–to–tail orientation of candidate telomeric retrotransposons (and the previously validated TAHRE and TART). Primer orientation to test head-to–tail positioning is represented above as cartoons in white and black. Gag (arrowhead) and RT (rectangle) domains are represented by lighter or darker shades, respectively. PCR-validated partial Gag and partial RT domains are represented as truncated symbols in *D. takahashii* and *D. ananassae.* **(B)** DNA-FISH or oligopainting with probes/paints cognate to TAHRE, TART, TARTAHRE and TR2 on polytene chromosomes from representative species. TART and TAHRE from D. melanogaster serve as positive controls. All insets show telomere hybridization exclusively except TARTARHE, which hybridized to both telomeric and non-telomeric locations (inset designated with a ‘*’).

The PCR-amplified/Sanger-sequenced domains typically share ~97.5% sequence identity to a given consensus (Table S3). For *D. takahashii* and *D. ananassae,* we successfully amplified the predicted partial TAHRE and TR2 RT domains, respectively, as well as *D. ananassae’s* partial TR2 Gag domain. We also confirmed head-to-tail orientation using primers that annealed to the 3’ ‘tail’ of one copy and the 5’ ‘head’ of another copy predicted to reside at the same telomere (Fig. 2A, Table S3). In *D. takahashii,* we successfully PCR-amplified an *in* silico-predicted junction between its partial TAHRE and a Helitron. Helitrons are DNA transposons that we discovered in subsequent experiments have colonized the telomeres of *D. takahashii’s* closest relative in our sampled species—*D. biarmipes* (see below). PCR- and Sanger sequencing-based validation suggests that virtually all consensus sequences represented in Fig. 1 correspond to actual jockey subclade elements found in their respective genomes and in an orientation stereotypical of telomere-specialized retrotransposons. Restriction to telomere ends, however, has only been demonstrated previously for the TAHRE and TART lineages.

To investigate chromosome localization of the newly defined elements, TR2 and TARTAHRE, we conducted DNA FISH or ‘oligopainting’ on polytene chromosomes in representative species. TAHRE/HeT-A and TART probes in *D. melanogaster* served as a positive control. Like the striking telomere-specialization of TAHRE/HeT-A and TART in *D. melanogaster,* the TR2 probe hybridized exclusively to telomere ends (Fig. 2B). However, TARTAHRE probe hybridization revealed *both* telomere localization and pericentromeric localization, as well as localization at other euchromatic sites (Fig. 2B). The promiscuous localization of TARTAHRE implicates either incomplete domestication or escape from domestication. Its nested phylogenetic position within a clade of telomere-specialized elements favors an innovation event leading to “escape” from the telomere (see Discussion).

### Recurrent turnover of telomere-specialized retrotransposons in the melanogaster species group

To infer telomeric retrotransposon lineage turnover across the melanogaster species group, we summarized presence/absence of these validated elements across the species tree (Fig. 3A, (Chen et al. 2014)). The TART lineage, well-known from *D. melanogaster,* is relatively young, emerging in the ancestor of the ‘subgroup’ between 10 and 15 million years ago. The TAHRE lineage is more ancient, emerging either along the lineage leading to the melanogaster species group or instead before the common ancestor of the melanogaster species group and *D. ananassae* (and subsequently lost along the *D. ananassae* lineage). TAHRE has been lost at least three times, along lineages leading to *D. rhopaloa, D. ficusphila* and *D. biarmipes. D. biarmipes’* closest relative on the tree, *D. takahashii,* encodes only a truncated TAHRE. These two species together suggest that TAHRE loss began in their common ancestor, though only in *D. biarmipes* is the loss event complete. The ‘generalist’ TARTARHE lineage appears in *D. rhopaloa* only, making it the only jockey subclade element restricted to a single species on the densely sampled, melanogaster species group tree. TR2 absence from *D. melanogaster* and its close relatives suggests the possibility that this lineage was functionally replaced by TAHRE and/or TART. Finally, we infer that TR2 was lost at least once along the lineage leading to the *D. takahashii/D. biarmipes* clade.

**Figure 3.**
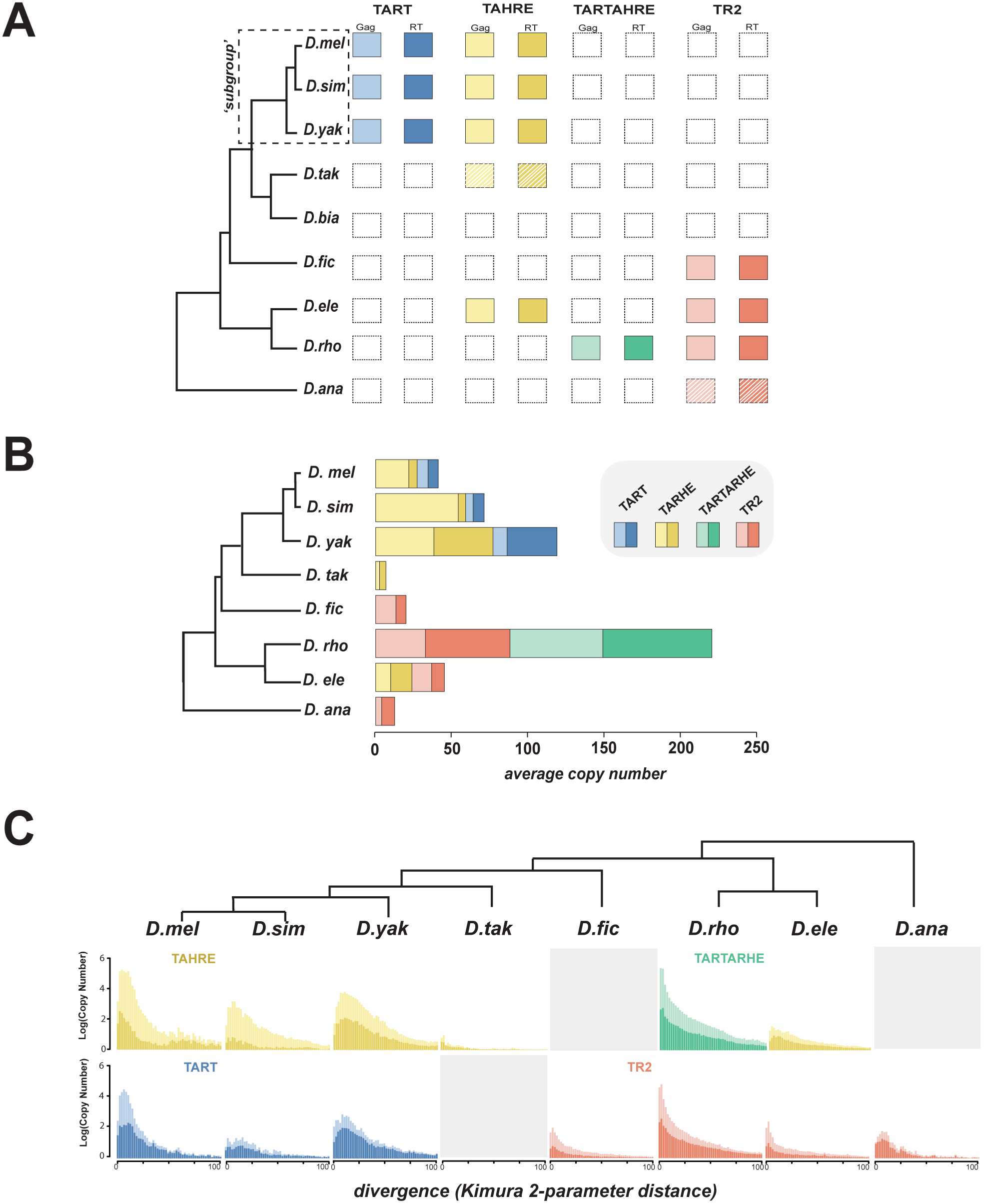
Element identity, copy number, and history across the melanogaster species group. **(A)** Presence/absence of telomere-localized elements across the melanogaster species group and *D. ananassae.* Each column represents a phylogenetically distinct lineage defined and validated in Fig. 1 and Fig. 2, respectively. Hatched lines delineate elements for which only a degraded version was recovered. Gag domains are represented by lighter shaded arrows and RT domains are shown as the darker shaded rectangles. **(B)** Estimated Gag (light) and RT (dark) copy number per species, calculated from the average read depth of a consensus sequence relative to the genome-wide estimate. **(C)** Repeat landscapes of telomere-specialized retrotransposons captured by Kimura 2-p distance between genomic reads with significant BLAST hits to the consensus (>90% identity). Copy number (consensus read # / genome-wide average read #) appears on the Y-axis and binned divergence classes appears on the X-axis. The bins closest to zero putatively represent the youngest classes. Gag (lighter shade), RT (darker shade).

While some lineages appear to rely on only one element, others harbor multiple retrotransposon lineages. The observation that *D. elegans,* for example, encodes both TAHRE and TR2, *D. rhopaloa* both TARTAHRE and TR2, and *D. melanogaster* both TART and TAHRE, rejects the null expectation that the retrotransposon lineage tree recapitulates the species tree. Instead, retrotransposon lineages are selectively retained and selectively lost across the species phylogeny, leading to retrotransposon tree-species tree discordance (Fig. 1 versus 3).

Overall, these data are consistent with diversification via gain and loss, both with and without replacement, of major retrotransposon lineages across three to 15 million years of evolution. The melanogaster species group is especially dynamic. We observe pervasive lineage-specific presence/absence of TR2, TARTAHRE, and TAHRE and even extreme cases of wholesale loss of jockey subclade, telomere-specialized elements.

### Expansions and contractions of DNA content derived from telomere-specialized elements

Retrotransposon expansions and contractions over time result in contemporary species-restriction of specific retrotransposon lineages; that is, progressively diminishing copy number precedes the wholesale loss of a retrotransposon lineage. To evaluate such bulk sequence changes across species, we mapped raw reads to a given domain consensus sequence and quantified read depth (Fig. 3B, Table S4). This copy number estimate serves also as a proxy for relative telomere length for all elements except TARTARHE, which localizes to both telomeric and non-telomeric sites. We note that the highly variable telomeric retrotransposon abundance within *D. melanogaster* (Wei et al. 2017) suggests that our single-genome estimates offer only a partial picture of divergence in bulk content.

We observe dramatic between-species differences in copy number, even for species that share common retrotransposon lineages. The degree and direction of these between-species differences were robust to multiple percent similarity thresholds to the consensus, minimizing the likelihood that we inadvertently underestimated copy number due to highly diverged variants (Table S4). qPCR on genomic DNA in cases of extreme copy number differences across species validated these computationally-generated estimates (Fig. S4). Across *melanogaster* subgroup species, telomeric retrotransposon content is 2-fold larger in *D. yakuba* compared to *D. melanogaster,* at least in the sequenced strains. Moreover, *D. melanogaster* and *D. simulans* telomeres are predominantly comprised of the TAHRE Gag (HeT-A), while *D. yakuba* encodes relatively more TART RT (Fig. 3B). TARTARHE in *D. rhopaloa* is especially abundant, an unsurprising discovery given its localization not only to chromosome ends but also to many other chromosomal locations (Fig. 2B). The telomere-restricted element, TR2, is also highly abundant in *D. rhopaloa* but depauperate in *D. ananassae. D. takahashii* too harbors low copy number of TAHRE Gag and RT domains. Notably, we were also unable to recover using *in silico* or PCR-based methods a single complete TAHRE domain in *D. takahashii* (we recovered only truncated versions of both domains), suggestive of ongoing loss of TAHRE-dependent telomere elongation. These data highlight the divergent jockey subclade content even across species that share a common retrotransposon lineage.

To further refine our snapshot of retrotransposon invasion and degeneration history, we generated frequency distributions of pairwise read divergence for each element in each species. If transposition events were recent, we expect the highest frequency classes to exhibit lowest divergence. Alternatively, an element that has undergone only an ancient episode of expansion exhibits an excess of high frequency reads with similar but elevated divergence from the consensus. Since fully functional Drosophila telomeres must constantly renew their telomere-specialized retrotransposon template, we expect actively transposing telomeric elements to have high similarity between consensus-mapping reads. Profiles of sequence divergence (estimated by Kimura 2-p distance) support this mechanism—virtually all distributions reveal an enrichment of reads diverging minimally or not at all (Fig. 3C). Indeed, most distributions are typical of recent transposition bursts characterized by a preponderance of highly similar sequences. We cannot formally rule out the homogenizing force of gene conversion to the excess of highly similar reads; however, the empirically verified rates of transposition in *D. melanogaster* (Kahn et al. 2000) suggests that gene conversion may contribute only moderately to the observed patterns. The distributions of *D. ananassae* TR2, *D. simulans* TART, and *D. elegans* TAHRE instead appear more uniform, consistent with ancient activity followed by mutation accumulation.

### Telomere-specialized retrotransposons are absent in D. biarmipes

Our data suggest that jockey subclade, telomere-specialized retrotransposons frequently occupy the melanogaster species group’s telomeres. However, these elements appear as full length in only a subset of the 10 species. To evaluate the possibility that *active* insertions by telomere-specialized retrotransposons can be lost completely, we exploited the most extreme case of telomeric retrotransposon degeneration uncovered by our pipeline. We detected no *D. biarmipes* telomere-specialized elements branching inside the monophyletic jockey subclade (Fig. 1). From these short-read data, we recovered instead a jockey family Gag (arrow, Fig. 1A). Hybridizing a FISH probe cognate to ‘jockey_1’ confirmed our phylogeny-based inference. Specifically, this computationally predicted jockey element localized across *D. biarmipes* chromosome arms but not to telomeres (Fig. S5), a typical, jockey family chromosomal distribution (Kaminker et al. 2002; Xie et al. 2013). To test the hypothesis that this exceptional species has lost jockey family retrotransposons at its chromosome ends, we performed long-read sequencing using the Pacific Biosciences (PacBio) SMRT sequencing platform. We predicted that upon assembling whole telomeres from *D. biarmipes,* we would discover a newly domesticated, telomere-specialized mobile element lineage unrelated to the jockey family, the wholesale loss of telomere-specialized mobile elements, or both.

Our long read-based assembly of the *D. biarmipes* genome revealed dramatically different chromosome end composition from the well-studied telomeres of *D. melanogaster* (Fig. 4A). Using the packages PBcR (Berlin et al. 2015) or DBG2OLC (Ye et al. 2016), we assembled *D. biarmipes* telomeres corresponding to *D. melanogaster’s* 2L, 2R, 3L, and 3R chromosomes. To determine composition in both species, we delineated *D. melanogaster* and *D. biarmipes* telomeric DNA as distal to the most terminally annotated, orthologous genes from *D. melanogaster* (the two species share a common karyotype (Deng et al. 2007)). Consistent with previous literature (Pardue and DeBaryshe 2011; Mason et al. 2016), *D. melanogaster* telomeres are dominated by telomere-specialized elements (HeT-A/TAHRE and TART) and satellite repeats (primarily “HETRP_DM’, Fig. 4A). In sharp contrast, the 50-200kb *D. biarmipes* telomeres harbor no jockey subclade retrotransposons, consistent with our short read-based pipeline results above. We uncovered instead DNA transposons called Helitrons, and to a lesser extent, Gypsy family LTR elements (Fig. 4A). We validated these *D. biarmipes* telomere assemblies in two ways. First, we observed 99% correspondence and no structural discrepancies between our assembled *D. biarmipes* telomeres and those of an independent, recently published hybrid assembly based on 454 and Nanopore reads (Fig. S6, (Miller et al. 2018)). Second, hybridization of FISH probes cognate to the telomeric Helitron and telomeric satellite sequence confirmed that we successfully assembled the termini of *D. biarmipes* chromosomes (Fig. 4B, insets).

**Figure 4.**
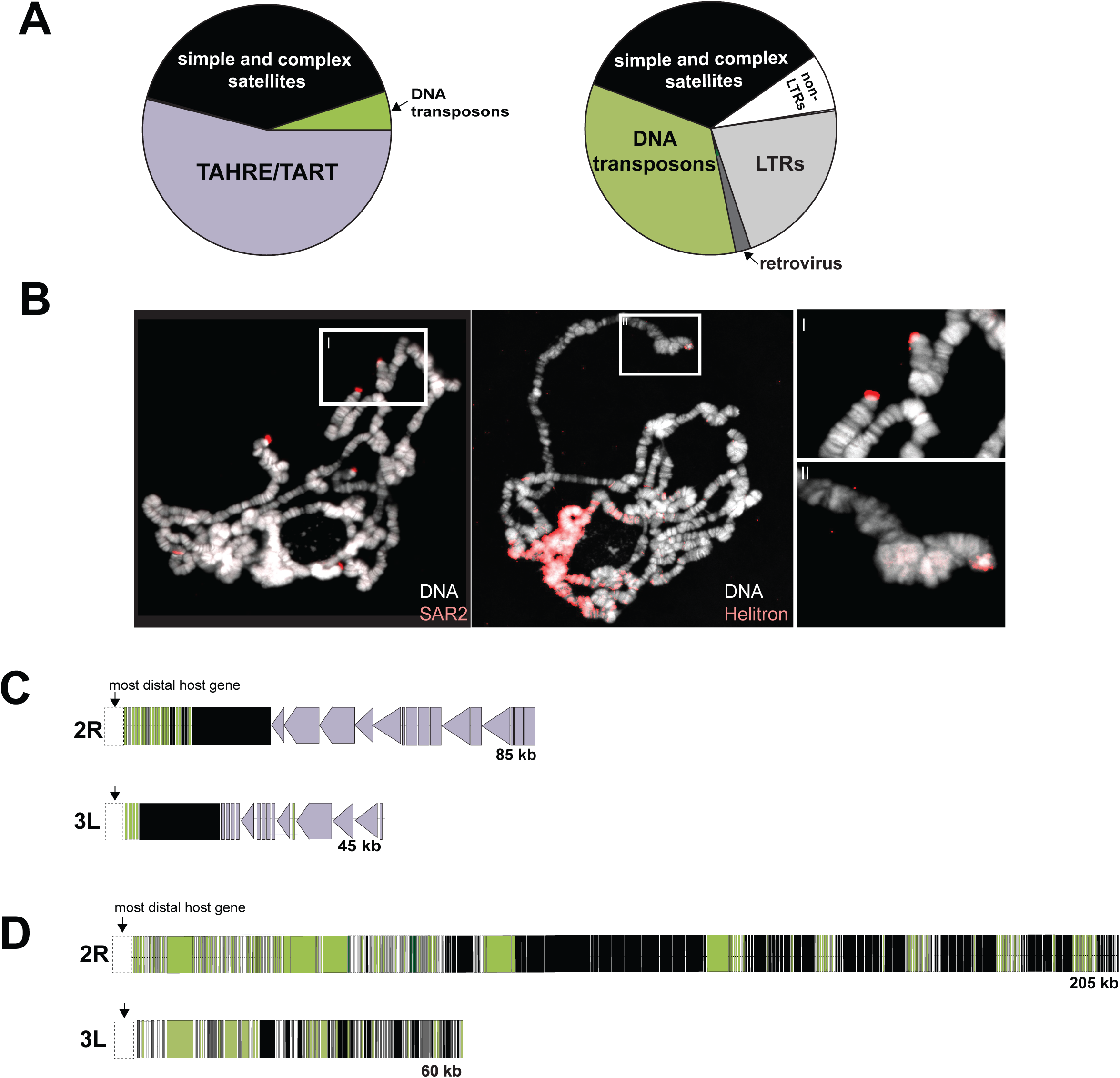
Collapse of the retrotransposon-based telomere elongation mechanism in *D. biarmipes*. **(A)** Composition of *D. melanogaster* and *D. biarmipes* chromosomes from the most distal, protein-coding gene and the terminal nucleotide (assembled from PacBio-generated long reads for both species) for Muller elements corresponding to 2L, 2R, 3L, and 3R. Purple corresponds to telomere-specialized, jockey family retrotransposons found in *D. melanogaster* (but not in *D. biarmipes).* **(B)** Fluorescent *in situ* hybridization of a SAR2 and Helitron probes to polytene chromosomes from *D. biarmipes.* Inset “I” and “II” shows telomere localization of SAR2 and Helitrons, respectively, on *D. biarmipes* polytene chromosomes. (C) Schematic representation of the long read-based assembly of *D. melanogaster* 2R and 3L telomeric DNA. Telomere-specialized, jockey-like elements (purple) are distal to a block of simple and complex satellites (black), consistent with previous reports. Triangles represent full-length elements and rectangles represent partially degenerated elements. (D) Schematic representation the of the long read-based assembly of 2R and 3L telomeres from *D. biarmipes* where simple and complex satellite DNA (black) is juxtaposed with primarily Helitron DNA transposons (green).

The rough equivalency of *D. melanogaster* and *D. biarmipes* telomeric DNA attributed to mobile elements and to satellite sequence suggests functional replacement of jockey subclade, telomere-specialized retrotransposons with other mobile element families. However, the integrity of these elements and their physical distribution challenges this inference. Starting at the most distally annotated host gene, *D. melanogaster* encodes long, uniform tracts of satellite DNA (primarily “HETRP-DM”, (Karpen and Spradling 1992)) followed by equivalently uniform tracts of the end-integrating TART and TAHRE (Fig. 4C, Table S5). Many of these elements are full-length and so presumably still active. Pervasive degradation instead dominates the Helitrons of *D. biarmipes* (Fig. S7). Moreover, these fragments are interspersed amidst satellite sequence (‘SAR’ and ‘SAR2’, Fig. 4D) and orientated randomly (Table S6), unlike the head-to-tail arrays of jockey subclade retrotransposons in *D. melanogaster.* All assembled *D. biarmipes* telomeres exhibit this pattern, despite varying in length from 50-200kb and varying in enrichment for transposable element family (Fig. 4D, Fig. S8, Table S6). Finally, the abundant Helitron signal at the chromocenter (Fig. 4B) and the patterns of divergence between copies (Fig. S9) highlights the similarity between *D. biarmipes* telomeres and its pericentric heterochromatin, consistent with ‘generalist’ mobile elements occupying a niche once-restricted to telomere-specialized elements.

Patterns of divergence between Helitron insertions and repeat structure of satellite sequence implicates an alternative lengthening mechanism in *D. biarmipes.* Using the uniquely long chromosome 2R as model, we observed that Helitron divergence is uniformly elevated between both nearby and distant insertions, whereas TAHRE Gag divergence is uniformly low (Fig. S10). We attribute this pattern of divergence to an ancient Helitron invasion followed by mutation accumulation. This pattern contrasts sharply with uniformly low divergence of TAHRE Gag from *D. melanogaster* (Fig. S10), likely driven by constant “refreshment” of end-inserting TAHRE Gag (“HeT-A”) elements. The absence of full-length Helitrons anywhere in our long-read *D. biarmipes* assembly (Fig.S7) suggests that these *D. biarmipes* telomeric Helitron fragments do not jump via Helitron mobilization machinery in *trans.* The *D. biarmipes* telomere tracts of repeats are instead reminiscent of Drosophila pericentromeric heterochromatin, which encodes long tracts of simple and complex satellites interspersed with dead TEs (Hoskins et al. 2007). *D. biarmipes* telomeres reveal the dramatic loss of a greater than 40 million-year old telomere elongation mechanism.

## Discussion

Virtually all eukaryotes rely on end-targeting and reverse transcription by telomerase to add DNA repeats to chromosome termini (Greider and Blackburn 1989; Zakian 1989; Blackburn 1991; Zakian 1996). The chromosome ends of *D. melanogaster* instead rely on telomere-specialized retrotransposons (Pardue and DeBaryshe 2011). These ‘domesticated’ mobile elements, too, end-target and reverse transcribe RNA into repetitive DNA at the most terminal base pairs, preserving unique genes several thousand nucleotides away. These alternative mechanisms differ not only in the molecular players that preserve chromosome ends but also in the rates of molecular evolution of telomeric DNA sequence. The telomerase-associated, nucleotide repeat unit composition changes slowly across eukaryotes (Meyne et al. 1989; Mason et al. 2016; Podlevsky and Chen 2016). Our investigation of the melanogaster species group reveals instead rapid diversification of the repeats charged with telomere length maintenance in Drosophila.

Building on previous discoveries of phylogenetically distinct, candidate telomeric jockey elements (Villasante et al. 2007), we report rapid turnover of major retrotransposon lineages across the melanogaster species group. Our comprehensive computational search, supported by experimental validation, revealed that an ancient TR2 lineage is retained in some species but sporadically lost and replaced by TAHRE, and/or TART in others. Notably, some species harbor a single retrotransposon lineage, like *D. ficusphila,* while others maintain multiple retrotransposon lineages, like *D. melanogaster* and *D. elegans.* Still others, like *D. biarmipes,* appear to lack telomere-restricted retrotransposons altogether. Consistent with a dynamic history of turnover, telomere-associated retrotransposon copy number also varies broadly. *D. rhopaloa,* for example, encodes 10x the TR2-derived sequence of *D. ananassae.* Appreciating that copy number also varies dramatically within species (Wei et al. 2017), we presume that between-species divergence documented here captures only a partial picture of copy number divergence. Together, these data highlight the remarkable diversity of telomere-specialized sequence—both identity and amount—across only 15 million years of Drosophila evolution.

Partially degenerated elements, rather than full length elements, dominate chromosome ends of several focal species, e.g., *D. takahashii* and *D. ananassae.* This diversity of telomere composition challenges the widely cited idea that active transposition is the primary mechanism of length regulation genus-wide (Casacuberta and Pardue 2003; Villasante et al. 2007). Active retrotransposons indeed populate the well-studied telomeres of *D. melanogaster*—the long read-based assembly of *D. melanogaster’s* telomeres ((Kim et al. 2014), Fig. 4C) supports decades of work defining this system (Biessmann et al. 1990; Mason and Biessmann 1995; Pardue and DeBaryshe 2003; Pardue and DeBaryshe 2011; Mason et al. 2016). Moreover, it is well-established that these active elements insert even at terminal deficiencies proximal to *D. melanogaster’s* telomeric retrotransposon array (Biessmann et al. 1990; Kahn et al. 2000). Finally, all Drosophila species investigated over the past several decades harbor telomere-specialized jockey subclade elements (Casacuberta and Pardue 2003; Berloco et al. 2005; Villasante et al. 2007). However, in contrast to *D. melanogaster,* species like *D. takahashii* appear to encode only partial copies of its telomeric retrotransposons (Fig. 2A, Fig. S11).

The predominance of only degenerated versions in this species raises the possibility that chromosome length maintenance depends on an alternative class of telomere-specialized mobile elements or instead, a non-mobile element-based mechanism. If newly domesticated mobile elements readily replace ancestral, telomere-specialized lineages, we would expect the species ostensibly lacking active jockey subclade elements—*D. biarmipes*—to encode such new recruits. The extreme terminal ends of *D. biarmipes* instead harbor AT-rich satellite “SAR” DNA (Mirkovitch et al. 1984; Kas and Laemmli 1992) at all telomeres as well as Helitron fragments (Kapitonov and Jurka 2007) and Gypsy-derived LTR fragments (Nefedova and Kim 2017) at variable frequencies across different telomeres. These non-autonomous elements are found both inside and outside the *D. biarmipes* telomeres (Fig. 4D) and found more typically outside the telomere in other Drosophila species (Kas and Laemmli 1992; Kapitonov and Jurka 2007; Nefedova and Kim 2017). Moreover, we found not one full-length Helitron in the *D. biarmipes* genome, rejecting the possibility of contemporary propagation using mobilization proteins in *trans* (Fig. S7). These data suggest that active transposition is not responsible for telomere length regulation, at least recently, in *D. biarmipes.* Intriguingly, Dias *et al.* (2015) previously reported dramatic expansion of a Helitron tandem repeat, DINE-TRI, along only the lineage leading to *D. biarmipes* and the lineage leading to *D. virilis/D. americana* relative to 25 other Drosophila genomes (Dias et al. 2015). Hybridization of DINE-TRI probes to *D. virilis* polytene chromosomes revealed striking localization to multiple telomeres, in addition to pericentromeric heterochromatin, reminiscent of the localization we detected in *D. biarmipes.* Like *D. biarmipes* telomeres, telomeric Helitrons in *D. virilis* represent only partial fragments rather than full-length, actively transposing copies (Dias et al. 2015). However, unlike *D. biarmipes, D. virilis* telomeres also encode jockey subclade retrotransposons predicted to maintain its chromosome length (Casacuberta and Pardue 2003).

In the absence of active telomeric transposition, how might *D. biarmipes* (and possibly *D. takahashii)* telomeres be maintained? *D. melanogaster* telomeres elongate not only by transposition but also by a recombination-based mechanism called ‘terminal gene conversion’ (Kahn et al. 2000; Melnikova and Georgiev 2002). This alternative lengthening pathway, utilized across eukaryotes alongside or instead of telomerase (McEachern and Haber 2006; Sakofsky and Malkova 2017; Sobinoff and Pickett 2017), relies on the shorter “broken” telomere resecting and invading a longer telomere. Second strand synthesis extends the chromosome end. This newly synthesized strand serves as the template for synthesis of its sister. In addition to terminal gene conversion, nonallelic recombination between two short telomeres could also lengthen deficient chromosomes, where the shorter product is ultimately removed by natural selection.

We predict that *D. biarmipes* depends on this alternative pathway. All studied, non-Drosophila species from the insect Order, Diptera, lack both telomerase-added repeats *and* telomere-specialized TEs and so are presumed to depend on terminal gene conversion (Mason et al. 2016). Specifically, Dipterans such as *Anopheles gambiae* (Biessmann et al. 1998), *Chironomous sp.* (Lopez et al. 1996; Rosen and Edstrom 2000), and *Rhynchosciara americana* (Lopez et al. 1996) encode only simple and/or complex satellites at chromosome ends. We, therefore, predict that Drosophila species lacking full length, telomere-specialized retrotransposons rely exclusively on this alternative elongation mechanism. Consistent with recombination shaping *D. biarmipes* telomere sequence evolution, we observe higher order repeat unit structure of telomeric SAR (Fig. S12). The concomitant proliferation of non-autonomous Helitrons at *D. biarmipes* telomeres may serve to replenish the repetitive sequence that mediates resected end homology searches and ultimately maintains the repair template. *D. biarmipes’* close relative, *D. takahashii,* likely represents a transitional state. In *D. takahashii,* we detected no evidence of a full-length TR2. Instead we recovered *in silico,* and then confirmed via PCR, at least one chimeric TR2-Helitron instance. The variable retention of a retrotransposon-based telomere elongation mechanism, together with the inferred ancestral Dipteran telomere state, helps us to contextualize what seemed initially like an aberration. Specifically, *D. biarmipes*’jockey element-poor, Helitron-rich telomeres may, in fact, represent a reversion to an ancestral, Dipteran-like state rather than a previously unseen innovation (Fig. 5).

**Figure 5.**
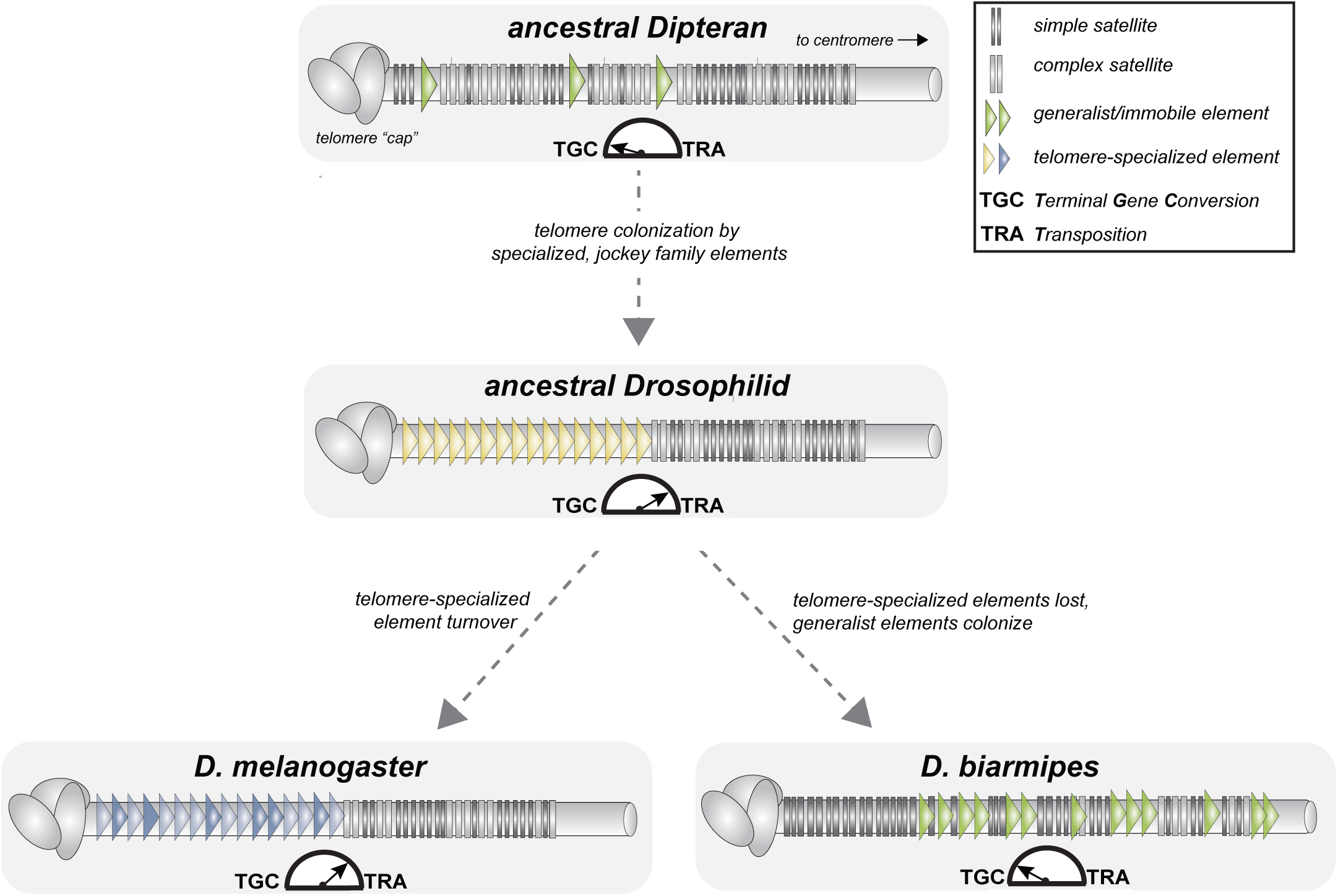
Model of telomere elongation mechanism evolution pre- and post- birth of Drosophilids. The Dipteran ancestor of Drosophila encodes neither telomerase nor telomere-specialized mobile elements. Instead, a recombination-based mechanism, possibly ‘terminal gene conversion’, lengthens the repetitive DNA (e.g., in mosquitos and midges). Exclusive chromosome-end insertions by a jockey family element becomes the primary, Drosophila-wide telomere elongation mechanism. Major jockey family lineages turn over across Drosophila species that retain this lengthening mechanism (bottom left). In species like D. biarmipes, the subsequent loss of telomere-specialized elements but presence of ‘generalist’ mobile elements illustrates how some Drosophila species may revert to the ancestral, predominantly recombination-based telomere lengthening mechanism (bottom right).

The discovery of “undomesticated” Helitrons at *D. takahashii* and *D. biarmipes* telomeres raises the possibility that some telomeric mobile elements act akin to ecological ‘facilitators’ rather than ‘mutualists’. Like a canopy tree inadvertently cultivating a favorable environment for shade-tolerant herbs, these immobile elements, by virtue of where they inserted during an ancient transposition burst, serve inadvertently the genome’s recombination-based telomere lengthening mechanism. Moreover, we cannot rule out the possibility that reverse transcription of satellite or Helitron RNA may also extend these telomeres (and other species’) by utilizing a non-telomeric retrotransposon’s reverse transcriptase in *trans* (Gorab 2003). We speculate that these ‘back up’ mechanisms reduce constraint on telomere-maintenance by active transposition, setting the stage for intra-genomic conflict to shape mobile element diversity across the melanogaster species group.

If intra-genomic conflict lies beneath this genomic symbiosis, who are the dueling parties? Compromised host fitness may ultimately select for expulsion of ‘disobedient’ retrotransposons that excessively lengthen telomeres or disrupt the euchromatic chromosome arms upon ‘escape’ from this chromosomal niche, as TARTAHRE appears to have done (host retrotransposon conflict). Indeed, evolutionary pressure to police these telomeric retrotransposons may shape the adaptive evolution detected at telomere length regulator proteins encoded by the host (Lee et al. 2017). Alternatively, transposable elements may compete among themselves for dominance within this genomic habitat (TE-TE conflict). More likely, both host genome and mobile element fitness together shape telomeric DNA diversity. Future work will define the genetic determinants and the molecular mechanisms shaping this ostensibly tenuous relationship between the host genome and its ever-evolving telomeric retrotransposons.

## Materials and Methods

### Species selection

To elucidate lineage turnover of Drosophila telomeric retrotransposons, we harnessed the ‘melanogaster species group’, which captures 15 million years of evolution (Chen et al. 2014). This slice of evolutionary time offers a previously unexplored scale of resolution: published investigations have targeted either short evolutionary distances from *D. melanogaster*—the melanogaster subgroup, where major telomeric retrotransposon lineages are conserved (Danilevskaya et al. 1998; Casacuberta and Pardue 2002; Berloco et al. 2005; Villasante et al. 2007)—or instead very long evolutionary distances from *D. melanogaster* (e.g., *D. ananassae, D. pseudoobscura),* where telomeric retrotransposon lineages are wildly diverged and never localized to chromosome ends (Villasante et al. 2007). The melanogaster species group spans an intermediate distance poised to capture the degradation of ancestral retrotransposons that maintain telomeres and the birth of new ones. *D. takahashii, D. ficusphila, D. biarmipes, D. elegans* and *D. rhopaloa* broadly sample this unexplored part of the Drosophila tree. To these species, we compare the melanogaster subgroup, including *D. simulans, D. sechellia, D. yakuba. D. ananassae* serves as a distant outgroup previously investigated by (Villasante et al. 2007).

### Drosophila genome sequence data used to define telomeric retrotransposons

Reference genomes assembled from short read sequences rarely include the repetitive DNA elements that accumulate at chromosome ends. We searched instead the publicly available raw sequence reads derived from 10 species (Table S7) sequenced with either Sanger (ABI 3700) and/or 454 (Genome Sequencer FLX). Although sequencing depth varies across these 10 species, we discovered jockey subclade, candidate telomere-specialized elements in even the lowest-coverage genome. The only exception was *D. biarmipes,* a species to which we ultimately subjected long-read sequencing (see below). Our single molecule-based genome assembly confirmed the absence of jockey subclade elements. For all other genomes, we detected no biased enrichment of jockey family lineages in genomes sequenced using one platform or another (Fig. S13). We also used publicly available Illumina-based short read data for consensus sequence correction (Table S7, see below).

### Identification of candidate telomeric retrotransposons from raw reads

We developed a custom method (Fig. S2) to search raw reads from 10 Drosophila species for the non-LTR retrotransposons related to jockey subclade of telomere-specialized elements. We designed this two-step method to identify phylogenetically distinct elements that maintain chromosome ends—both active and degraded copies. Our initial query included all previously characterized, telomeric-specialized retrotransposons from the melanogaster subgroup *(D. melanogaster, D. simulans, D. sechellia, D. yakuba)* as well as those uncharacterized elements only predicted to maintain telomere ends in more distant species (Table S1). To ensure that our search was exhaustive, we included even retrotransposons for which telomeric localization or head-to-tail orientations had not been confirmed (Table S1, references therein). We also included the reference jockey element from *D. melanogaster* (Repbase, www.girinst.org/Repbase/). These input sequences served as the query for a tBLASTn search of a given species’ raw read database (Table S7). We retained any read that shared 80% sequence identity with any one of the query elements. We then *de novo* assembled this subset of raw reads into consensus sequences (i.e., two or more reads assembled by the Geneious assembler: Medium Sensitivity/Fast parameters). We aligned each consensus sequence to the query sequences plus the reference jockey element from *D. melanogaster* using MAFFT (k =2, Gap penalty=1.53, Offset=0.123). For each alignment of either the Gag or RT domain, we built a phylogenetic tree using FastTree (GTR+CAT, (Price et al. 2009)) and determined if the focal consensus sequence branched as an ingroup or an outgroup (outside the jockey element). We retained only ingroup consensus sequences for subsequent analysis (482 out of 4,583 consensus sequences).

We repeated this pipeline by inputting the 482 consensus sequences from round one (length mean= 4143, standard deviation= 1656) as a new query in a BLASTn search of the raw reads from each species. This second iteration enriched our pool substantially: we obtained ~2 million total reads from all 10 species in this round compared to 60,000 from the first round (Table S8). We filtered uninformative reads by mapping the two million reads to the 482 consensus sequences sited above (Geneious mapper: Medium Sensitivity/Medium; Iterative Fine Tuning = 5). We refined consensus accuracy from the pool of 754,230 reads retained with filtering. Again, we *de novo* assembled the pool of retained reads, which generated 62,383 new consensus sequences. We repeated the FastTree sorting method and retained only consensus sequences that branched internally to the jockey clade, leaving us with 3,112 consensus sequences (length mean= 1531, standard deviation = 732).

We next generated full-length Gag and RT domains from our consensus sequences. We first translated the nucleotide sequences using “six-pack” (Rice et al. 2000) and aligned using MAFFT to known telomere-specialized retrotransposon domains (parameters: -k = 2, Gap penalty=1.53, Offset=0.123, Table S1, (Katoh and Standley 2013)). We then parsed the Gag and RT consensus sequences based on its branch position *(i.e.* TAHRE-like, TART-like, TR2-like, jockey-like) from the previous FastTree sorting step. Guided by the alignments, we removed frameshifts and unalignable sequence. We discarded entire consensus sequences harboring less than 80% identity over 80% of its alignment length to any other consensus within the subset (Wicker et al. 2007). For each subset of consensus sequences per species, we generated final consensus sequences using majority rule. We also used Repbase to infer consensus identity for all discarded sequences described above. These sequences represented mostly jockey elements (Table S2) and also several highly degraded, telomere-associated elements juxtaposed with DNA unrelated to the jockey family. We conducted a final refinement step for each complete Gag and RT domain by mapping raw reads to our set of telomeric retrotransposon candidates. These raw reads came from Illumina-based short read sequence databases generated independently of the Sanger and 454 reads used to build the consensus sequences (Table S7). This step successfully called many sites previously designated ambiguous based on majority rule. In addition, we elongated the domain boundaries using reads that spanned the consensus and sequence outside the domain. Again, we used majority rule to infer the final sequence. We repeated cycles of mapping and extension up to 10 times until we detected a full-length open reading frame and, in most cases, the sequence spanning Gag and RT for a given element. All scripts are available upon request.

### Phylogenetic tree building of final consensus sequences

We built Gag- and RT- based phylogenetic trees of the retained consensus non-LTR retrotransposons using MrBayes 3.2.6 (Ronquist et al. 2012), implemented in Geneious (Kearse et al. 2012). We determined the amino acid substitution model using ProTest 3.4.2 (Darriba et al. 2011). We performed MCMC analysis over 1,100,000 generations. We estimated convergence of the joint posterior probability density each 200 generations with a 25% burn-in of the sampled chains. We stopped the analysis once the maximum standard deviation of split frequencies was < 0.001. The *D. melanogaster* reference jockey element served as the outgroup. We built phylogenies of Gag and RT domains defined by our pipeline using the majority-rule consensus sequences only (Fig. 1). We also built Gag and RT trees that included all previously described telomeric retrotransposon consensus within the melanogaster subgroup and all previously predicted candidate telomeric elements outside of the melanogaster species group to contextualize our results in the previous literature (Table S1, Fig. S3).

### Validation of computationally-defined candidate telomeric retrotransposons

Our custom pipeline detected many species-restricted, non-LTR retrotransposons that form a monophyletic clade with previously described telomere-specialized elements. These phylogenetically distinct, intact open reading frames are supported by overlapping reads along each sequence and high coverage (>27X on average). However, assemblies of short reads mapping to repetitive sequence and majority rule-based consensus sequences may include computational artifacts that do not represent sequences that truly exist in a given genome.

#### PCR validation

We used PCR to validate 1) domain consensus sequences and 2) head-to-tail tandem array orientation stereotypical of telomere-lengthening retrotransposons. Specifically, we designed primers (Table S9) from our consensus sequences and performed PCR on genomic DNA (Table S10) prepared from 10 females from each species (ArchivePure™, 5 PRIME). We gel-extracted and Sanger-sequenced (Genewiz, South Plainfield, NJ) the PCR products of the expected size (QIAquick^^®^^ Gel Extraction kit, Table S3).

#### Cytogenetic validation

PCR-validated elements may be phylogenetically related to telomere-specialized elements but not necessarily telomere-restricted. We used fluorescent *in situ* hybridization (FISH) probes and/or ‘Oligopaints’ (Beliveau et al. 2015) to determine whether the computationally predicted, PCR-validated retrotransposons localize to chromosome ends. Specifically, we hybridized to polytene chromosomes FISH probes cognate to HeT-A, TART (both *D. melanogaster* controls), and TARTAHRE *(D. rhopaloa)* as well as Oligopaints cognate to TR2 *(D. elegans,* (Table S9)).

We also evaluated whether a jockey family element from *D. biarmipes,* a species that lacks HeT-A, TART, TARTAHRE, and TR2, localized to telomeres using FISH (previously referred to HTR0 (Villasante et al. 2007)). Finally, we hybridized Helitron and SAR2 sequence-derived FISH probes to *D. biarmipes* polytene chromosomes to validate our long read-based assembly of its chromosome ends (Table S9).

We first dissected salivary glands from third-instar larvae in a PBS-Tween 0.1% solution. We then transferred salivary glands into fresh 45% acetic acid and 2% paraformaldehyde for one minute on a coverslip treated with Sigmacote^®^ (Sigma-Aldrich). We lowered Poly-L-lysine (Sigma-Aldrich) treated slides onto the coverslip and spread the polytene chromosomes by tapping the slide with a rubber hammer. We flash-froze the slides in liquid nitrogen and transferred into 100% ethanol (−20°C) for 10 minutes. To prepare chromosomes for probe hybridization, we washed twice in PBS-Tween 0.1% for 5 minutes (min) and then transferred slides in successive pre-hybridization solutions (5 minutes in 2X SSC plus 0.1% Tween, 5 minutes in 2X SSC plus 0.1% Tween and 50% formamide, 2.5 min at 92°C in 2X SSC plus 0.1% Tween and 50% formamide, and then 20 min at 60 °C in 2X SSC plus 0.1% Tween and 50% formamide).

We labeled DNA FISH probes either with DIG dUTP (Sigma-Aldrich, HeT-A, TART, TARTAHRE) or used Oligopaints (TR2, Table S9). We diluted probes (15μg/ml for PCR DIG probes and 2μg/ml for Oligopaints) in a hybridization buffer (50% formamide, 25% Dextran Sulfate, 10μg of RNase A and 12.5% of ddH2O), denatured probes for 2.5 min at 95 °C and then incubated on ice for 2 min. We performed hybridization overnight at 50°C for PCR-DIG probes and 37°C for Oligopaints. We removed unbound probes with two 10 min washes in 2X SSC at 60 °C and three 10 min washes in PBS at room temperature. We stained PCR-DIG probes with anti-Dioxigenin-AP Fab fragments (Sigma-Aldrich), diluted 1:250 in PBST+BSA 3%, 1-hour incubation) and then anti-Sheep IgG (H+L) Alexa Fluor 555 conjugated secondary antibody (Invitrogen™, diluted 1:500 in PBST+BSA 3%, 1-hour incubation). For Oligopaints, we followed the (Beliveau et al. 2015) protocol for secondary incubation. We visualized hybridization experiments on a Leica SP8 confocal microscope.

### Relative telomeric retrotransposon copy number and estimates of within-genome diversity

To estimate the relative abundance of our validated telomeric retrotransposons in a given species genome, we mapped raw reads (Sanger or 454) to both our consensus sequences and the most recent genome assembly (Table S7) using SMALT (http://sourceforge.net/projects/smalt/) with varying parameters (k: length of the hashed word index, s: the sampling distance between successive words, and y: the percent identity allowing a word to match). We selected the parameters that maximized coverage for both genome and TE consensus (index: -k 20, -s 13; map: -y 0.9). To infer retrotransposon copy number, we divided a given retrotransposon average coverage by the genome-wide average coverage. Previous reports suggested that a non-trivial fraction of telomere-specialized retrotransposons are partially degenerated (Mason and Biessmann 1995). We indeed detected widespread degradation in all sampled genomes. To account for retrotransposon sequence represented by these reads encoding partially degraded sequence, we also conducted a BLAST-guided approach to capture all reads harboring >90% identity to the consensus sequence. We trimmed these reads from their non-telomeric specific sequence and re-calculated normalized coverage with these trimmed reads. We report the latter analysis. In addition, we estimated divergence between these filtered reads and retrotransposon consensus in order to infer retrotransposon invasion history. We aligned in PHYLIP a given consensus sequence and all significant, >90% identity reads. After visual inspection, we estimated the rates of transitions and transversions from alignment files and calculated Kimura 2-parameter distances (Kimura 1980).

#### Validation of copy number estimates

We evaluated our computational predictions of copy number variation using qPCR on genomic DNA prepared from 10 females per species (ArchivePure™, 5 PRIME). We designed primers to amplify the region of highest mean coverage (estimated from the BLAST analysis above) and used the ΔΔCq method to quantitate abundance relative to a single copy gene (rp49).

### Single molecule sequencing, assembly, and validation of *D. biarmipes*

#### DNA preparation for single molecule sequencing

We detected no evidence of retrotransposons related to TAHRE, TART, TARTARHE, or TR2 in the (short) raw reads of *D. biarmipes.* To define the set of telomeric genetic elements found in this unusual species, we conducted single-molecule sequencing using the Pacific Biosciences platform. We prepared high quality DNA using Qiagen Blood and Cell culture DNA MIDI Kit according to (Chakraborty et al. 2018) with some modifications (Qiagen). Specifically, we collected 150mg of tissue from 200 zero to two-day old, whole females. We held the females in empty vials for two hours prior flash-freezing in liquid nitrogen and grinding to a fine powder. We transferred the sample into 9.5ml of buffer G2 with addition of 38μl of RNase-A (100mg/ml) and 250μl of protease (0.75AU). We homogenized the sample and then incubated at 50°C overnight with gentle shaking. The next day, we centrifuged the sample at 5000 x g for 10 min at 4°C. We decanted the supernatant into a fresh 15ml tube, vortexed, and applied it to the anion exchange column. We washed the column and precipitated genomic DNA with 0.7 volume of isopropanol resuspended in Tris buffer (pH 8) and stored at 4°C overnight. Genewiz (South Plainfield, NJ) prepared a 20 kb PacBio SMRTbell library with Blue Pippin size selection per manufacturer’s instruction.

#### Assembling D. biarmipes genome with long reads

We built both PacBio-only assemblies and PacBio/Illumina hybrid assemblies. We used the PBcR pipeline (Celera 8.3) to assemble PacBio reads only. First, we executed a self-correction step (‘PBcR-MHAP’, k = 14, merSize=16, sketches = 1024, coverage = 25), which generated a pre-assembly (N50= 480 kb; longest contig = 4.3 Mb). We passed the Celera assembler (CA 8.3) the longest 25X coverage corrected contigs, using parameters optimized for genomes >100Mb (Berlin et al. 2015). The N50 and longest contig both increased to a final 718 kb and 9.1 Mb, respectively. We generated a hybrid assembly of our PacBio reads and the publicly available Illumina short reads from *D. biarmipes* (Table S7) using the software package DGB2OLC (Ye et al. 2016), following parameters described in (Chakraborty et al. 2018). This hybrid assembly increased the N50 and longest contig to 2.5Mb and 9.2Mb, respectively. Both assemblies and raw reads have been deposited at NCBI under XXXXXX.

#### Defining telomeric DNA from D. biarmipes

To extract telomeric DNA sequence from the *D. biarmipes* PBcR assembly, we relied on the well-annotated, full-length chromosome arms of *D. melanogaster* (Release r6.17). Starting with a query of terminal *D. melanogaster* genes (~20 genes per chromosome arm), we used BLASTn to identify *D. biarmipes* contigs encoding terminal sequence *(D. biarmipes* and *D. melanogaster* share a common karyotype (Deng et al. 2007)). We identified exactly four *D. biarmipes* contigs with homology to *D. melanogaster* 2L, 2R, 3L and 3R chromosome ends. We next used these four contigs as queries in a BLASTn search of the *D. biarmipes* assembly to identify any additional contigs with homology to the terminal sequence. We identified only a single, 29kb contig corresponding to 3R, which overlapped the original 3R contig over 15kb (>99% identity). This new contig augmented the 3R telomeric sequence by an additional 14kb of distal sequence. To further validate our assembly of four *D. biarmipes* telomeres, we compared them to a second hybrid assembly generated independently from Nanopore and Illumina reads (Miller et al. 2018). In no case did we observe in Miller *et al.* contigs that extend beyond the most distal sequence from our assembly (see Fig. S6).

#### Analysis of telomeric DNA sequence in D. biarmipes and D. melanogaster

For both our *D. biarmipes* telomeric contigs and those defined using the same methods for *D. melanogaster* (Kim et al. 2014), we annotated the sequence distal to the most terminal gene using RepeatMasker (Smit, AFA, Hubley, R & Green, P. *RepeatMasker Open-4.0).* From this annotation, we plotted TE family abundance (Fig. S8) and then focused our analyses on the two highest repeat classes, the Helitron 2N DNA transposon and the SAR2 complex satellite repeat. We extracted the repeats from annotated contigs and aligned them to their respective consensus. From these PHYLIP alignments, we calculated Kimura 2-parameter distance. We summarized distance estimates on heatmaps according to their position along telomere terminal sequences (Fig S10). We also annotated higher order repeat structure of individual SAR2 variants from PHYLIP alignments by ordering SAR2 repeat according to their position along telomeres (Fig S12). Finally, we compared telomeric Helitrons to peri-centromeric (those sharing a contig with most proximal genes) and euchromatic Helitrons by parsing copies from our PacBio assembly using the Helitron 2N consensus as a BLASTn query (>80% identity threshold). Using the methods described above, we calculated Kimura 2-p distance of copies derived from telomeric, pericentromeric, and euchromatic regions as above (Fig S9).

## Data Access

All consensus sequences and PCR-amplified elements can be found in Table S2 and S3, respectively. Our PacBio-generated long reads for *D. biarmipes* were deposited at NCBI under SRA XXXXX. Genome assemblies will be deposited at www.flybase.org.

## Author Contributions

M.T.L and B.S. designed the experiments. B.S. performed the bioinformatics, the experiments, and the analyses. S.C.N. designed the Oligopaints. M.T.L and B.S. wrote the manuscript.

## ACKNOWLEDGEMENTS

The authors thank C. Leek for assistance with PCR-based validation experiments and G. Lee and S. Zanders for their comments on earlier versions of the manuscript. This work was supported by an NIH NIGMS R00GM107351 and an NIH NIGMS R35GM124684 to MTL.

